# *Rpi-amr3* confers resistance to multiple *Phytophthora* species by recognizing a conserved RXLR effector

**DOI:** 10.1101/2021.06.10.447899

**Authors:** Xiao Lin, Andrea Olave-Achury, Robert Heal, Kamil Witek, Hari S. Karki, Tianqiao Song, Chih-hang Wu, Hiroaki Adachi, Sophien Kamoun, Vivianne G. A. A. Vleeshouwers, Jonathan D. G. Jones

## Abstract

Diverse pathogens from the genus *Phytophthora* cause disease and reduce yields in many crop plants. Although many *Resistance to Phytophthora infestans* (*Rpi*) genes effective against potato late blight have been cloned, few have been cloned against other *Phytophthora* species. Most *Rpi* genes encode nucleotide-binding domain, leucine-rich repeat-containing (NLR) proteins, that recognize RXLR effectors. However, whether NLR proteins can recognize RXLR effectors from multiple different *Phytophthora* pathogens has rarely been investigated. Here, we report the effector AVRamr3 from *P. infestans* that is recognized by Rpi-amr3 from *S. americanum*. We show here that AVRamr3 is broadly conserved in many different *Phytophthora* species, and that recognition of AVRamr3 homologs enables resistance against multiple *Phytophthora* pathogens, including *P. parasitica* and *P. palmivora*. Our findings suggest a novel path to identifying *R* genes against important plant pathogens.

## Introduction

Species in the oomycete genus *Phytophthora* cause many devastating plant diseases. For example, *P. infestans, P. parasitica, P. cactorum, P. ramorum, P. sojae, P. palmivora* and *P. megakarya* cause disease on potato and tomato, tobacco, strawberry, oak, soybean and cacao, respectively (Kamoun et al., 2015).

Plant immunity involves detection of pathogen-derived molecules by either cell-surface pattern recognition immune receptors (PRRs) or intracellular nucleotide-binding domain, leucine-rich repeat containing (NLR) immune receptors, that activate either pattern-triggered immunity (PTI) or effector-triggered immunity (ETI), respectively (Jones and Dangl, 2006). So far, many *Resistance to P. infestans* (*Rpi*) genes were cloned from wild *Solanum* species which confer resistance against potato late blight (Vleeshouwers et al., 2011). Many *R* genes against *P. sojae* (*Rps)* have also been mapped in different soybean accessions, and some were cloned (Sahoo et al., 2017). In tobacco, the black shank resistance genes *Phl, Php* and *Ph* were genetically mapped but not yet cloned; these confer race-specific resistance to *P. parasitica* (aka *P. nicotianae*) isolates (Gallup and Shew, 2010; Bao et al., 2019). For *P. palmivora*, some resistant cacao (*Theobroma cacao*) accessions were identified, but no dominant *R* genes have been defined or cloned (Thevenin et al., 2012). In summary, very few *R* genes against other *Phytophthora* pathogens have been cloned.

*Solanum americanum* and *S. nigrum* are highly resistant to *P. infestans* (Witek et al., 2016; Witek et al., 2021). Two *Rpi* genes of coiled-coil (CC) type, *Rpi-amr3* and *Rpi-amr1*, were cloned from different *S. americanum* accessions, and both confer broad-spectrum late blight resistance in cultivated potatoes (Witek et al., 2016; Witek et al., 2021).

In oomycetes, Rpi proteins typically recognize RXLR (Arg-X-Leu-Arg, X represents any amino acid)-EER (Glu-Glu-Arg) effectors that are secreted into plant cells (Rehmany et al., 2005; Wang et al., 2019). Many *Avirulence* (*Avr*) genes encoding recognized effectors from *Phytophthora* species have been identified and they are often fast-evolving and lineage-specific molecules (Jiang et al., 2008). Recently, AVRamr1 (PITG_07569), the recognized effector of Rpi-amr1 was identified by a long read and cDNA pathogen enrichment sequencing (PenSeq) approach (Lin et al., 2020a). Surprisingly, AVRamr1 homologs were identified from *P. parasitica* and *P. cactorum* genomes and both are recognized by all Rpi-amr1 variants (Witek et al., 2021). Similarly, AVR3a-like effectors were found in different *Phytophthora* species, including *P. capsici* and *P. sojae*, and the recognition of AVR3a homologs correlates with *P. capsici* or *P. sojae* resistance in *Nicotiana* species and soybean (Shan et al., 2004; Vega-Arreguín et al., 2014), also AVRblb2 homologs from *P. andina* and *P. mirabilis* trigger HR with Rpi-blb (Oliva et al., 2015). Remarkably, a single N336Y mutation in R3a expands its recognition specificity to *P. capsici* AVR3a homolog (Segretin et al., 2014). These reports raise intriguing questions. Could RXLR effectors be widely conserved molecules among different *Phytophthora* species? Could these effectors be recognized by the same plant receptor? Of particular interest, could this effector recognition capacity enable disease resistance?

Here we identified AVRamr3, a novel AVR protein from *P. infestans*, by screening an RXLR effector library. AVRamr3 is a broadly conserved effector found in many different *Phytophthora* species. Strikingly, the recognition of AVRamr3 not only enables resistance to *P. infestans*, but also to other economically important *Phytophthora* pathogens including *P. parasitica* and *P. palmivora*. We also show functional *Rpi-amr3* genes are widely distributed among *S. americanum* and *S. nigrum* accessions, together with the previously defined *Rpi-amr1* genes, they might underpin the “non-host” resistance of these species against *P. infestans*.

## Results

### *Avramr3* encodes a conserved RXLR-WY effector protein

To identify the effector recognized by Rpi-amr3, we screened an RXLR effector library (Rietman, 2011; Lin *et al*., 2020) by *Agrobacterium tumefaciens*-mediated co-expression with Rpi-amr3 in *Nicotiana benthamiana*. By screening ∼150 RXLR effectors, we found PITG_21190 specifically induces hypersensitive response (HR) with *Rpi-amr3* (Fig. 1a), and concluded PITG_21190 is *Avramr3. Avramr3* encodes a 339-aa protein with a signal peptide followed by RXLR, EER motifs and four predicted WY motifs (Win et al., 2012) (Fig. 1b). In *P. infestans* T30-4, the expression of *Avramr3* is low during infection, however *Avramr3* upregulated in 3928A and US23 in 2-3 days after infection (Cooke et al., 2012; Lin et al., 2020a).

**Figure 1.**
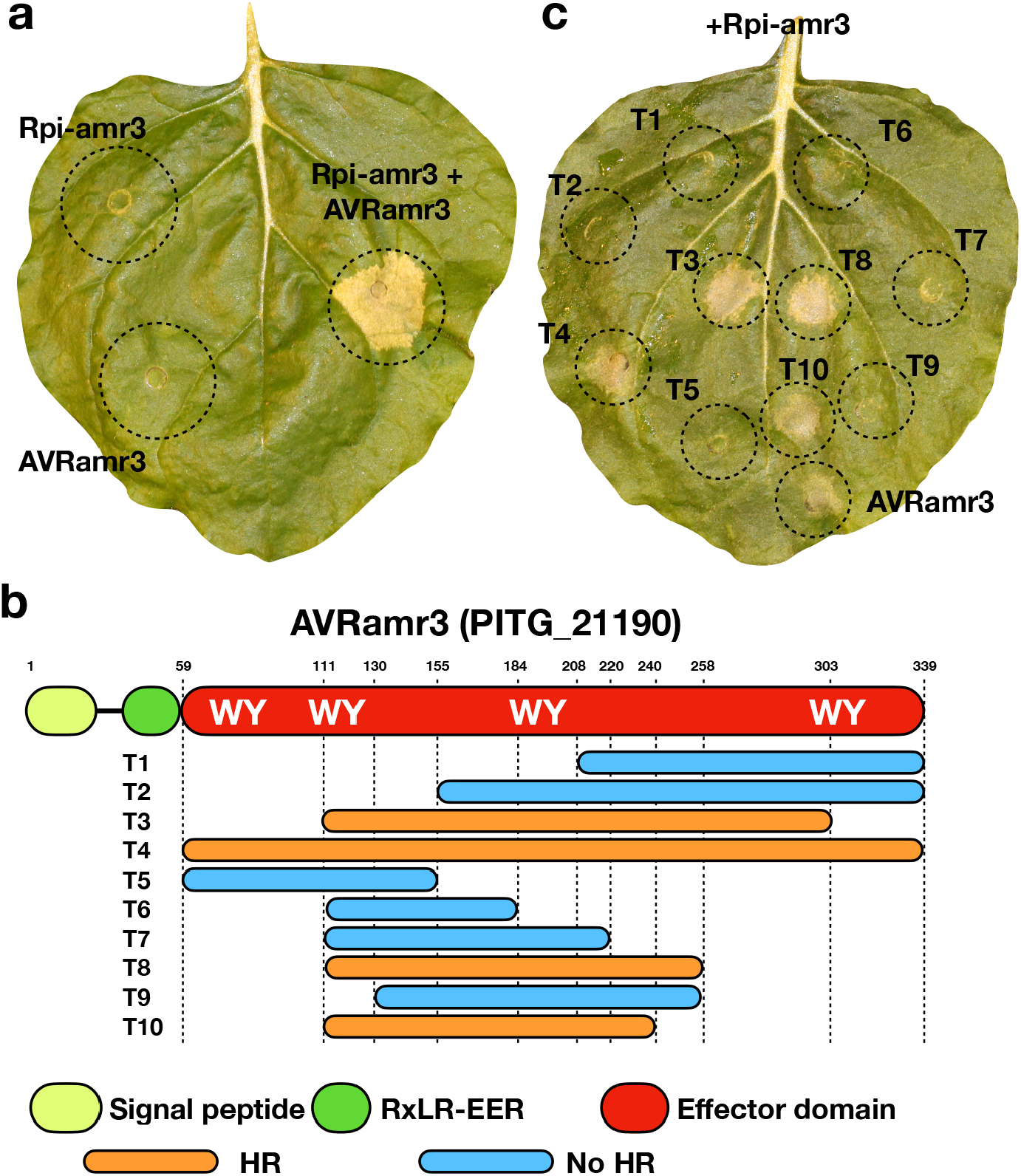
AVRamr3 is the recognized effector for Rpi-amr3. **(a)**. Co-expression of Rpi-amr3::GFP and AVRamr3::HIS-FLAG trigger cell death on *N. benthamiana*, but expression of Rpi-amr3::GFP or AVRarm3::HIS-FLAG individually do not induce cell death. The photo was taken 3 days after infiltration, *Agrobacterium* strain GV3101(pMP90) carrying Rpi-amr3:GFP or AVRamr3::HIS-FLAG constructs were used in this experiment. OD_600_=0.5. Three biological replicates were performed with same results. **(b)**. Cartoon of AVRamr3 (PITG_21190), a protein with 339 amino acids with a signal peptide (lemon), RXLR-EER motif (green), and an effector domain (red) with four predicted WY motifs (Details are shown in Fig. S4). T1-T10 indicate the AVRamr3 truncations used in HR assays. Those that induce HR after co-expression with Rpi-amr3 are marked by orange bars, otherwise by blue. **(c)**. Co-expression of Rpi-amr3::GFP and AVRamr3 truncations, all truncations are tagged with C-terminal HIS-FLAG tag. T3, T4, T8 and T10 trigger cell death when co-expressed with Rpi-amr3, but not T1, T2, T5, T6, T7 and T9. Full-length AVRamr3::HIS-FLAG was used as control. OD_600_=0.5. Three biological replicates were performed with same results.

Many RXLR effectors are fast-evolving, multiple-member family proteins with extensive sequence polymorphism, such as the *Avr2* and *Avrblb2* families (Gilroy et al., 2011; Oliva et al., 2015). To study the sequence polymorphism of *Avramr3*, seventeen additional *Avramr3* homologs from eleven isolates were identified from published databases (KR_1, 3928A, EC1, 6_A1 and US23) (Lee et al., 2020; Lin et al., 2020a) or cloned by PCR (EC1, Katshaar, Pi14538, Pi88069, Pi99183 and Pi99177) (Fig. S1). The sequence alignment shows *Avramr3* is a highly conserved RXLR effector among *P. infestans* isolates (Fig. S1).

To find the domain responsible for recognition by Rpi-amr3, ten truncated *Avramr3* fragments were cloned (T1 to T10, Fig. 1b, Fig S2) in an expression vector, and transiently co-expressed with *Rpi-amr3* in *N. benthamiana*. We found four AVRamr3 truncations (T3, T4, T8 and T10) can be recognized by Rpi-amr3. T10 (111-240 aa) which carries the 2^nd^ and 3^rd^ WY motifs is the minimal region to be recognized by Rpi-amr3 but not the adjacent T9 protein (130-258 aa) (Fig. 1b and 1c). This suggests these 130 amino-acids of AVRamr3 T10 are sufficient for recognition by Rpi-amr3 and initiation of HR.

### Rpi-amr3 is dependent on the helper NLRs NRC2, NRC3 and NRC4

In Solanaceae, the functionality of many CC-NLR proteins requires helper NLR proteins of the NRC class (Wu et al., 2017). To test if *Rpi-amr3* is NRC-dependent, we co-expressed *Rpi-amr3* and *Avramr3* in NRC knockout *N. benthamiana* lines (nrc2/3_1.3.1, nrc4_185.9.1.3, nrc2/3/4_210.4.3)(Adachi et al., 2019; Wu et al., 2020; Witek et al., 2021). As with wild type *N. benthamiana*, we found HR on the nrc2/3_1.3.1 and nrc4_185.9.1.3 knockout lines, but not the nrc2/3/4_210.4.3 knockout line. Similarly, only nrc2/3/4_210.4.3 knockout lines show susceptibility to *P. infestans* after *Rpi-amr3* transient expression (Fig. S3). Therefore, these data suggest both *Rpi-amr3*-mediated effector recognition and resistance require NRC2, NRC3 or NRC4.

### Rpi-amr3 associates with AVRamr3 *in planta*

To date, most Rpi proteins recognize their cognate effectors in an indirect manner, except RB and IPI-O effectors (Chen et al., 2012; Kourelis and van der Hoorn, 2018). To test the interaction between Rpi-amr3 and AVRamr3, Rpi-amr3::HA and AVRamr3::HIS-FLAG epitope-tagged constructs were generated and transiently co-expressed in nrc2/3/4 knockout *N. benthamiana* leaves to avoid cell death. Protein was then extracted and bi-directional co-immunoprecipitation (Co-IP) was performed. These co-IPs indicate that Rpi-amr3 associates with AVRamr3 bidirectionally (Fig. 2a). We also tested their interaction using a split-luciferase assay. Rpi-amr3::Cluc and AVRamr3::Nluc constructs were transiently expressed in the nrc2/3/4 knockout *N. benthamiana*. Luciferase signal was only detected when Rpi-amr3::Cluc and AVRamr3::Nluc were co-expressed. It suggests Rpi-amr3 physically associates with AVRamr3 *in-planta* (Fig. 2b), but not in negative controls. Our data therefore are consistent with direct interaction of Rpi-amr3 and AVRamr3 proteins, though do not exclude the possible involvement of additional proteins.

**Figure 2.**
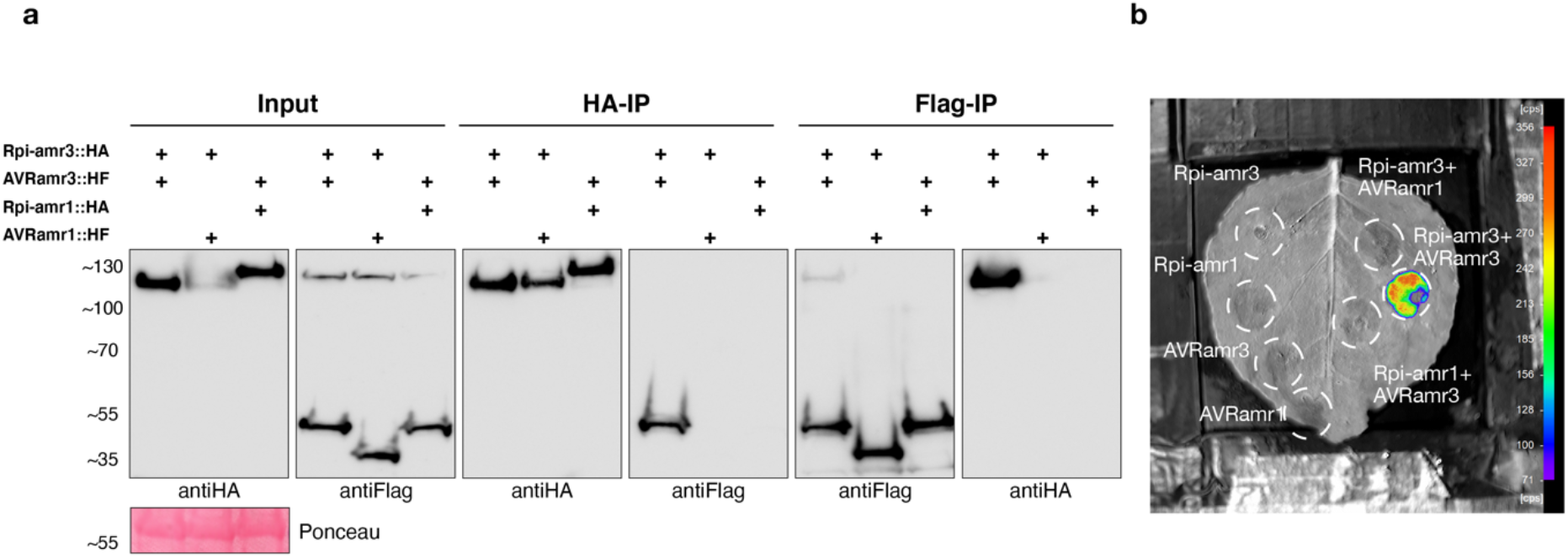
Rpi-amr3 directly interacts with AVRamr3. **(a)**. Rpi-amr3::HA and AVRamr3::HIS-FLAG constructs were used for bidirectional co-immunoprecitation experiment, with Rpi-amr1-HA and AVRamr1::HIS-FLAG used as control. After HA pull down of Rpi-amr3::HA or Rpi-amr1::HA, only AVRamr3::HIS-FLAG is associated with Rpi-amr3::HA. After Flag pull down of AVRamr3::HIS-FLAG or AVRamr1-HIS-FLAG, only Rpi-amr3::HA is associated with AVRamr3::HIS-FLAG. *Agrobacterium* strain GV3101(pMP90) carrying different constructs were used for transiently expression in nrc2/3/4 knockout *Nicotiana benthamiana* line (210.4.3) to abolish the cell death phenotype. OD_600_=0.5. Three biological replicates were performed with same results. **(b)**. Rpi-amr3::Cluc and AVRamr3::Nluc constructs were used to test their interaction *in planta*, Rpi-amr1::Cluc and AVRamr1::Nluc were used as controls. The luciferase signal can only be detected upon Rpi-amr3::Cluc and AVRamr3::Nluc co-expression.

### *Avramr3* orthologs occur in multiple *Phytophthora* species

To study the evolution of *Avramr3* in *Phytophthora* species, we searched for *Avramr3* homologs from published *Phytophthora* and *Hyaloperonospora arabidopsidis* (Hpa) genomes. Surprisingly, we found *Avramr3* homologs in many *Phytophthora* genomes, including *P. parasitica, P. cactorum, P. palmivora, P. pluvialis, P. megakarya, P. lichii, P. ramorum, P. lateralis, P. sojae. P. capsici, P. cinnamomi*, and in *H. arabidopsidis*. Most of the *Avramr3* homologs are located at a syntenic locus (Fig. 3a). Notably, the *P. infestans Avramr3*-containing contig was not fully assembled; sequences are missing on the 5’ side of *Avramr3* (Fig. 3a). The protein alignment of the thirteen AVRamr3 homologs is shown in Fig S4.

**Figure 3.**
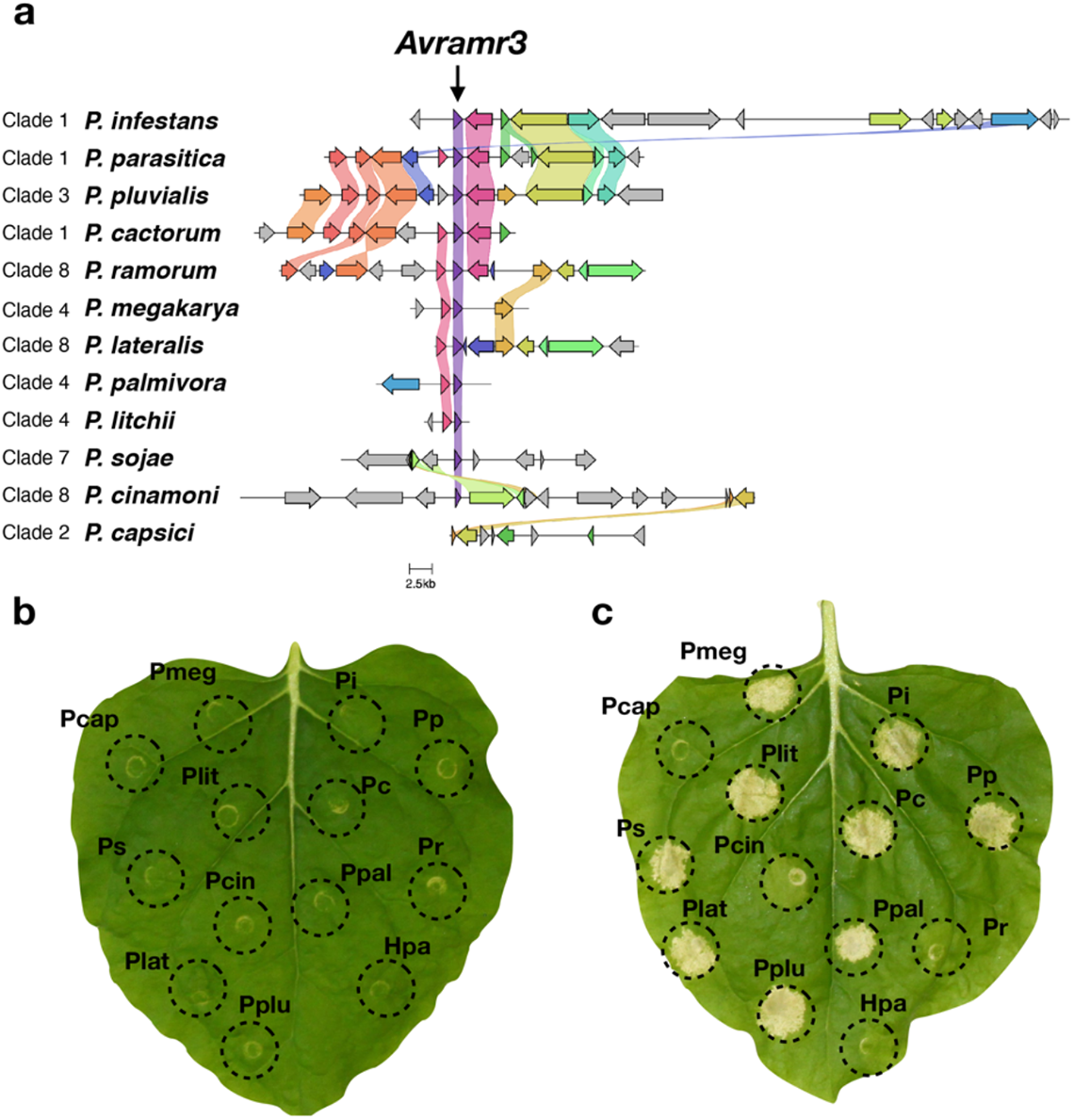
AVRamr3 is a conserved effector among different *Phytophthora* species. (a). The synteny map of *Avramr3* loci from twelve different *Phytophthora* genomes. The *Avramr3* loci were extracted from different genomes, annotated by the gene prediction tool in EumicrobeDB, then analyzed and visualized by Clinker. *Avramr3* homologs are shown by purple triangles and indicated by a black arrow, the flanking genes with homology are represented by the corresponding colours. The *Phytophthora* clades are adapted from *Phytophthora* database (Rahman et al., 2014). (b). Expression of AVRamr3 homologs with HIS-FLAG tag alone does not trigger cell death on *Nicotiana benthamiana. Agrobacterium* strain GV3101(pMP90) carring different constructs were used in this experiment. OD_600_=0.5. Three biological replicates were performed with same results. (c). Co-expression of AVRamr3 homologs with Rpi-amr3::GFP in *N. benthamiana*. The AVRamr3 homologs from *Phytophthora infestans* (Pi), *P. parasitica* (Pp), *P*.*cactorum* (Pc), *P. palmivora* (Ppal), *P. megakarya* (Pmeg), *P. litchi* (Plit), *P. sojae* (Ps), *P. lateralis* (Plat) and *P. pluvialis* (Pplu) induce cell death after co-expression with Rpi-amr3::GFP, but not AVRamr3 homologs from *P. ramorum* (Pr), *P. capsici* (Pcap) and *Hyaloperonospora arabidopsidis* (Hpa). The AVRamr3 homolog from *P. cinnamomi* (Pcin) shows an intermediate cell death. *Agrobacterium* strain GV3101(pMP90) carrying different constructs were used in this experiment. OD_600_=0.5. Three biological replicates were performed with same results.

To test if those AVRamr3 homologs from different *Phytophthora* species are also recognized by Rpi-amr3, we synthesized and cloned them into an expression vector with the 35S promoter, and performed transient expression assays in *N. benthamiana*. Expressing the effectors alone does not trigger HR in *N. benthamiana* (Fig. 3b), but AVRamr3 homologs from *P. parasitica, P. cactorum, P. palmivora, P. megakarya, P. lichii, P. sojae, P. lateralis* and *P. pluvialis* can induce HR when co-expressed with Rpi-amr3. The AVRamr3 homolog from *P. cinnamomi* triggers an intermediate HR, and the AVRamr3 homologs from *P. ramorum, P. capsici*, and *H. arabidopsidis* (Fig 3c) do not trigger Rpi-amr3-dependent HR.

To test if particularly conserved amino-acids of AVRamr3 are responsible for the Rpi-amr3 recognition, we mutated eight conserved amino-acid on the AVRamr3 T10 region (Figure S4). However, all tested mutants are still recognized by *Rpi-amr3* (Figure S5). This result indicates the recognition specificity might not be determined by any single amino acid on AVRamr3, but rather by its overall structure.

To test if other recognized AVRamr3 homologs also directly interact with Rpi-amr3, we performed co-immunoprecipitation and split-luciferase assays in nrc2/3/4_210.4.3 knockout lines. We found all the recognized AVRamr3 homologs associate with Rpi-amr3 by co-immunoprecipitation, though with varied affinity. Two unrecognized AVRamr3 homologs from *P. capsici* and *H. arabidopsidis* do not associate with Rpi-amr3. However, two unrecognized or weakly recognized AVRamr3 homologs from *P. ramorum* and *P. cinnamomi* also associate with Rpi-amr3, and the unrecognized AVRamr3-T9 truncation shows a weak association (Figure S6). In contrast, the output of split-luciferase assay is fully consistent with the HR assay (Figure S7). Our data indicating that an *in-planta* receptor-ligand interaction is necessary but might not be sufficient for the activation of Rpi-amr3 and triggering of HR.

### Rpi-amr3 confers resistance to multiple P. parasitica and P. palmivora> strains in N. benthamiana

*Rpi-amr3* was previously reported to confer resistance against potato late blight caused by *P. infestans* (Witek et al., 2016). Its broad effector recognition capacity suggested *Rpi-amr3* might confer resistance against additional *Phytophthora* pathogens.

To test this hypothesis, we generated *Rpi-amr3* stable transformed *N. benthamiana* lines. Two homozygous T2 lines #13.3 and #16.5 were verified to confer *P. infestans* resistance and evaluated for *P. parasitica* and *P. palmivora* resistance. Both these pathogens have a wide host range, including the model plant *N. benthamiana*.

Six *P. parasitica* isolates (R0, R1, 310, 666, 329 and 721) were tested on *N. benthamiana* carrying *Rpi-amr3*, and on wild type *N. benthamiana* plants as negative control. A suspension of zoospores was used for root inoculation (Material and Methods). We found both *N. benthamiana* – *Rpi-amr3* lines resist three *P. parasitica* isolates R1, 666 and 721, but are susceptible to R0, 310 and 329 (Fig 4). In summary, *Rpi-amr3* confers resistance against three out of six tested *P. parasitica* isolates in *N. benthamiana*.

**Figure 4.**
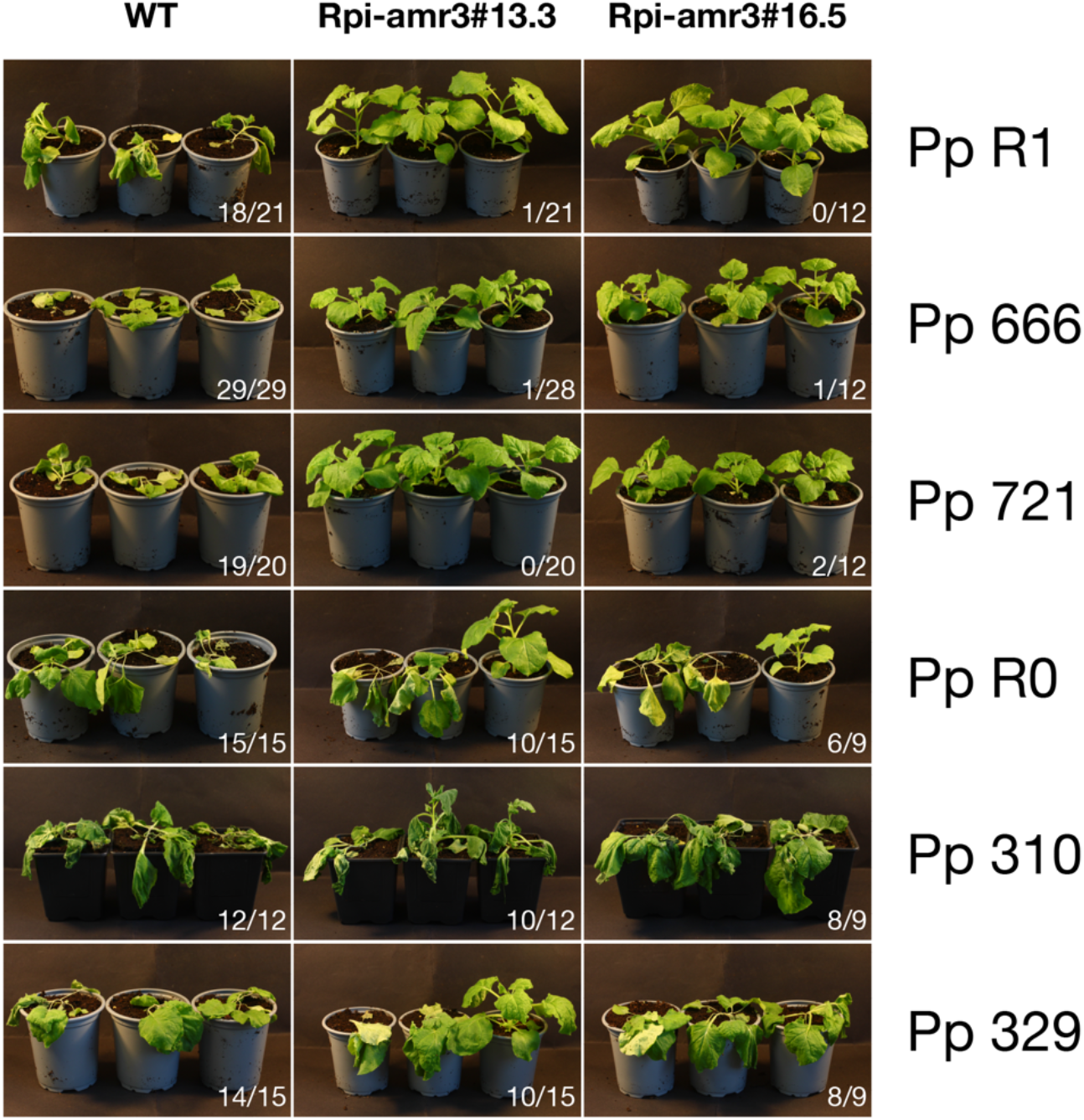
Root inoculation of six *Phytophthora parasitica* isolates on *Rpi-amr3* transgenic *Nicotiana benthamiana* lines. Representative photos for the *P. parasitica* root inoculation tests are shown. Two homozygous *N. benthamiana* - *Rpi-amr3* lines #13.3 and #16.5 were used in this experiment. Wild type *N. benthamiana* plants were used as control. Six *P. parasitica* isolates were used for root inoculation, *Rpi-amr3* confers resistance against R1, 666 and 721, but not R0, 310 and 329. 3-4 weeks *N. benthamiana* were used for the root inoculation, 3 plants/line were used for each experiment and as least three biological replicates were performed with similar results The numbers indicate susceptible plants/total tested plants.

The *PpAvramr3* homologs from the six *P. parasitica* isolates were PCR amplified, sub-cloned and sequenced. *PpAvramr3* homologs were identified from R0, R1 and 310, 666 and 721, but not from 329 (Fig S8). These data suggest the presence of recognized AVRamr3 homologs from the *Phytophthora* pathogens is necessary but not sufficient to induce Rpi-amr3 mediated resistance.

Furthermore, we tested another broad host range *Phytophthora* pathogen, *P. palmivora*, which causes major losses on many tropical tree crops like papaya, mango, cacao, coconut and palm tree. We tested seven *P. palmivora* isolates on the two *Rpi-amr3* transgenic *N. benthamiana* lines by root inoculation, and wild type *N. benthamiana* was used as a control. We found *Rpi-amr3* confers resistance to three out of seven tested *P. palmivora* isolates, including 7551, 7547, 7545, but not to 3914, 7548. For two other isolates 0113 and 3738, inconsistent results were obtained from the two *Rpi-amr3* transgenic lines (Fig 5). To verify the presence of *Avramr3* homologs in these tested *P. palmivora* isolates, we PCR amplified the *Avramr3* homologs from genomic DNA of the seven *P. palmivora* isolates. All the tested *P. palmivora* carry *PpalAvramr3* variants (Figure S9). Taken together, *Rpi-amr3* confers resistance to at least 3/7 tested *P. palmivora* isolates in the root inoculation assay.

**Figure 5.**
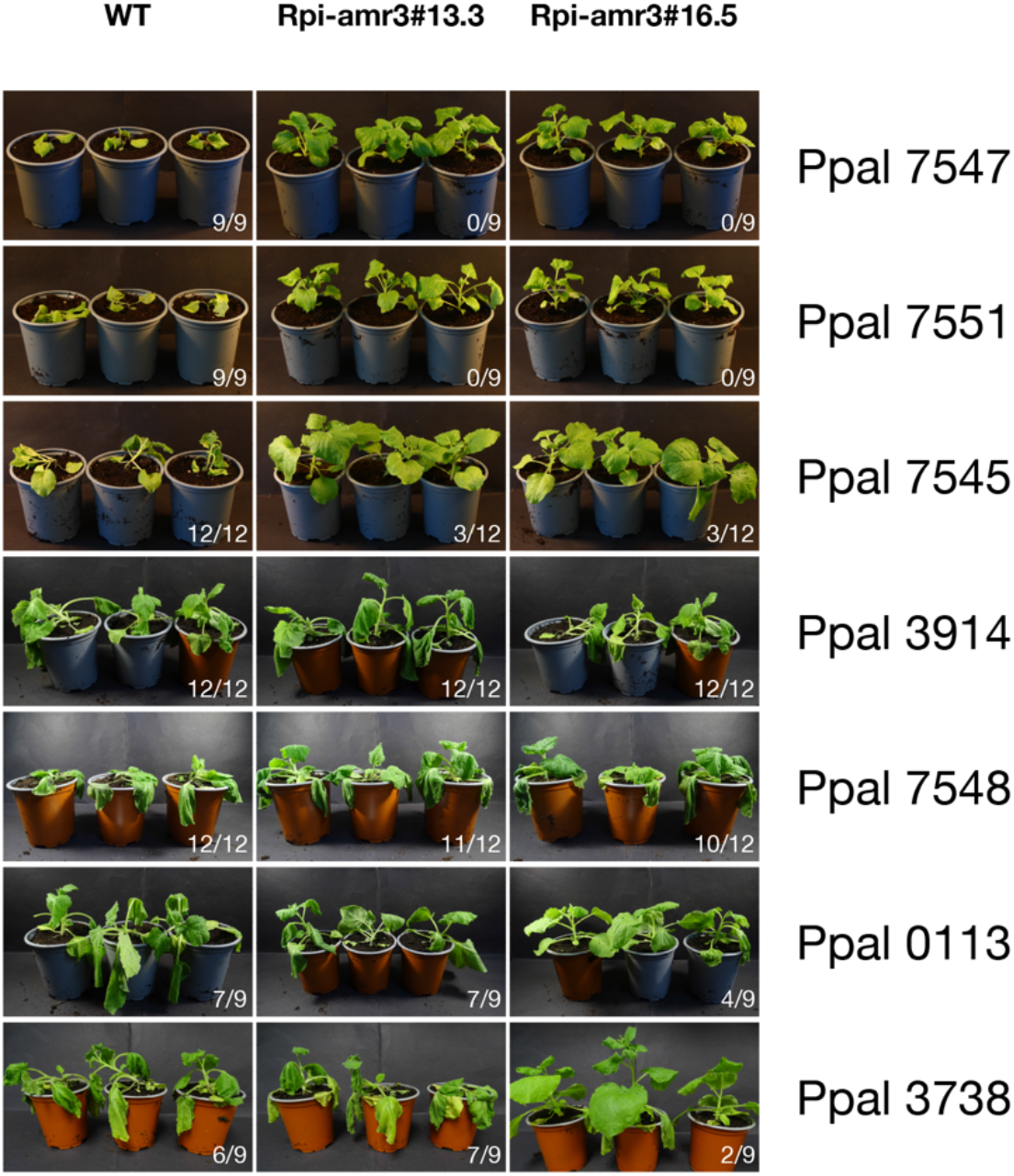
Root inoculation of 7 *Phytophthora palmivora* isolates on *Rpi-amr3* transgenic *Nicotiana benthamiana* lines. Two homozygous *N. benthamiana* - *Rpi-amr3* lines #13.3 and #16.5 were used in this experiment, wild type *N. benthamiana* were used as control. Seven *P. parasitica* isolates were used for root inoculation, *Rpi-amr3* confer resistance against isolates 7547, 7551 and 7545, but not 3914, 7548. For isolates 0113 and 3738, we obtained some variable results for the two transgenic lines. 3-4 old weeks *N. benthamiana* were used for the root inoculation, 3 plants/line were used for each experiment and three or more biological replicates were performed with similar results.

### Rpi-amr3 is widely distributed in S. americanum and S. nigrum

Though susceptible accessions can be identified in detached leaf assays, most *S. americanum* and *S. nigrum* accessions show complete resistance in the field to *P. infestans*. Previously, many functional *Rpi-amr1* alleles were cloned from different *S. americanum* and *S. nigrum* accessions (Witek et al., 2021).

The identification of AVRamr3 allows us to investigate the distribution of *Rpi-amr*3 from all *S. americanum* and *S. nigrum* accessions. In total, 54 *S. americanum* accessions and 26 *S. nigrum* accessions were tested by agro-infiltration with AVRamr3 for detecting functional *Rpi-amr3*. We found 43/54 tested *S. americanum* accessions show HR after AVRamr3 agro-infiltration (Fig 6a). Similarly, 21/26 tested *S. nigrum* accessions recognize AVRamr3 (Fig 6b).

**Figure 6.**
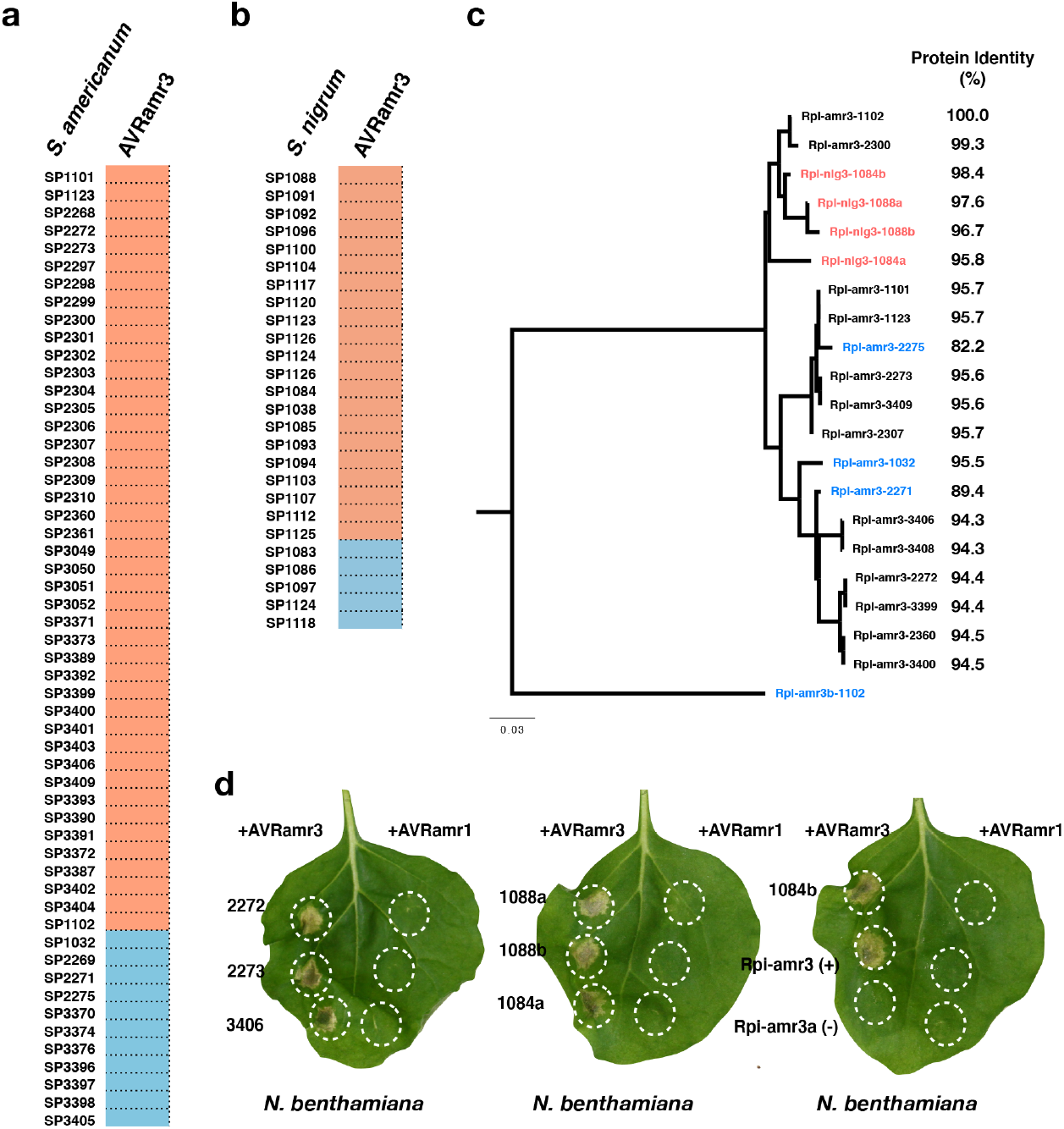
Screen for AVRamr3 recognition on *S. americanum* and *S. nigrum* accessions. **(a)**. 54 *S. americanum* accessions were screened with *Agrobacterium* strain GV3101(pMP90) carrying 35S::AVRamr3. The accessions with cell death upon agro-infiltration are marked by red, otherwise blue. **(b)**. 26 *S. nigrum* accessions were screened with *Agrobacterium* strain GV3101(pMP90) carrying 35S::AVRamr3. The accessions with cell death upon agro-infiltration are marked by red, otherwise blue. **(c)**. Maximum likelihood (ML) tree of Rpi-amr3 and Rpi-nig3 proteins was made by iqtree with GTT+G4 model. The *Rpi-amr3* homologs from *S. americanum* were extracted from PacBio RenSeq assemblies (Witek et al., 2021). The four *Rpi-nig3* genes were PCR amplified from *S. nigrum* accession SP1088 and SP1084 (red). The non-functional *Rpi-amr3* homologs were marked by blue. Rpi-amr3b from SP1102 is a paralogue of Rpi-amr3, which was used as an outgroup of the phylogenetic analysis. The scale bar indicates the number of amino acid substitutions per site. The protein identities of each homolog compared to Rpi-amr3 (Rpi-amr3-1102) are shown by %. **(d)**. Selected Rpi-amr3 homologs were cloned from three *S. americanum accessions* SP2272, SP2273, SP3406. Four Rpi-nig3 homologs were cloned from *S. nigrum* accessions SP1088 and SP1084, and co-expressed with AVRamr3 or AVRamr1 (negative control). All of them can recognize AVRamr3 in the transient assay but not AVRamr1.

To further investigate the sequence polymorphism of *Rpi-amr3* from different accessions, the *Rpi-amr3* homologs from 14 accessions were extracted from PacBio RenSeq dataset (Witek et al., 2021), including eleven accessions (SP1123, SP2272, SP2273, SP2307, SP2360, SP3399, SP3400, SP1101, SP3406 and SP3409) which respond to AVRamr3 and three accessions (SP1032, SP2271 and SP2275) that do not respond to AVRamr3.

To test the functionality of *Rpi-amr3* from *S. americanum* and *S. nigrum, Rpi-amr3* homologs were PCR amplified from gDNA of three *S. americanum* accessions SP2272, SP2273 and SP3406, and from gDNA of two *S. nigrum* accessions SP1088 and SP1084. *Rpi-amr3* alleles (*Rpi-nig3* hereafter) were amplified from each of these two *S. nigrum* accessions and cloned into an expression vector with 35S promoter. We found all the seven *Rpi-amr3*/*Rpi-nig3* genes can recognize AVRamr3 in transient assays (Fig 6d), but not the negative control AVRamr1. Compared to Rpi-amr3 from SP1102, the amino-acid identity ranges from 82.2% to 95.7% (Fig 6c). Premature stop codons were found in *Rpi-amr3* homologs from SP2271 and SP2275 (Figure S10), which result in loss of *Rpi-amr3* function.

Taken together, these data suggest *Rpi-amr3* gene is widely distributed in diploid *S. americanum* and hexaploid *S. nigrum*, and contributes to their resistance to *P. infestans* and perhaps other *Phytophthora* pathogens.

## Discussion

In this study, by screening an RXLR effector library of *Phytophthora infestans*, we identified and characterized a novel effector AVRamr3 (PITG_21190) that is recognized by the NLR protein Rpi-amr3 of *S. americanum*. AVRamr3 is very conserved among all tested *P. infestans* isolates, and AVRamr3 homologs were identified in twelve additional *Phytophthora* and *Hyaloperonospora arabidopsidis* genomes. These homologs are located in a syntenic region (Fig. 3a). Surprisingly, we found 9/13 tested AVRamr3 homologs can be recognized by Rpi-amr3 leading to HR in *N. benthamiana*. This finding suggests AVRamr3 is an essential effector among *Phytophthora* species, though its virulence function has yet to be determined.

According to the “zigzagzig” model of plant immunity (Jones and Dangl, 2006), the surface immune receptors like receptor-like proteins (RLPs) and receptor-like kinases (RLKs) perceive relatively conserved microbe-associated molecular patterns (MAMPs) and induce pattern-triggered immunity (PTI). Intracellular nucleotide-binding and leucine-rich repeat immune receptors (NLRs) recognize fast-evolving and lineage-specific effectors and activate effector-triggered immunity (ETI). Therefore, PTI was believed to confer broader-spectrum resistance compared to ETI. Indeed, many RLPs/RLKs recognize conserved ligands and /or confer broad-spectrum resistance, such as FLS2, EFR, RLP23, RXEG1 and ELR (Zipfel et al., 2006; Albert et al., 2015; Du et al., 2015; Wang et al., 2018). Remarkably, EFR from *Arabidopsis thaliana* enhances resistance against a range of bacterial pathogens in different crop plants, like tomato, orange and apple (Lacombe et al., 2010; Mitre et al., 2021; Piazza et al., 2021). However, apoplastic effectors can also be fast-evolving proteins, like the SCR74 family in *P. infestans* (Liu et al., 2005; Lin et al., 2020b), or *Cladosporium fulvum* AVR2, AVR4 and AVR9 (Joosten et al., 1994; Van den Ackerveken et al., 1994; Luderer et al., 2002; Westerink et al., 2004). On the other hand, MAMP-like cytoplasmic effectors/effector epitopes have been reported. For example, Sw-5b from tomato confers broad-spectrum tospovirus resistance by recognizing a conserved, 21-amino acid epitope NSm^21^ which derives from the viral movement protein NSm (Zhu et al., 2017). A recent functional pan-genome study revealed the ETI landscape of *A. thaliana* and *Pseudomonas syringae*; some *P. syringae* effectors are widely conserved. Similarly, the ETI mediated by two conserved NLRs CAR1 and ZAR1 confers resistance to 94.7% *P. syringae* strains (Laflamme et al., 2020). These observations, as well as our finding on AVRamr3 and Rpi-amr3, all support the view that pathogen molecules recognized by NLRs can also be relatively invariant and conserved, and might contribute to broad-spectrum pathogen resistance.

To test this hypothesis, we established a *N. benthamiana* root inoculation system by using stable *Rpi-amr3* transgenic plants, and tested *P. parasitica* and *P. palmivora* which have a broad host range, and cause dramatic yield losses of many crops from different families (Meng et al., 2014; Ali et al., 2017). Importantly, we found *Rpi-amr3* does confer resistance against some *P. parasitica* and *P. palmivora* isolates. This is the first report of cloned *R* genes against *P. parasitica* and *P. palmivora* (Kourelis et al., 2021). Additionally, it is noteworthy that, in nature, many *Phytophthora* pathogens can co-inoculate the host and interspecific hybridization might occur; for example, *P. andina* was proposed to have emerged through hybridization of *P. infestans* and an unknown *Phytophthora* species (Goss et al., 2011). Natural hybrids of *P. parasitica* and *P. cactorum* were also found on infected loquat trees (Hurtado-Gonzales et al., 2017). An R protein that provides protection against both foliar and root *Phytophthora* pathogens of different species would be extremely valuable. However, some *Rpi-amr3* breaking *P. parasitica* and *P. palmivora* strains were also identified in our study, although most of them carry the recognized AVRamr3 homologs. This might be caused by silencing of the recognized effector gene like *Avrvnt1* to avoid the recognition by Rpi-vnt1, or presence of other suppressors or regulators like Avrcap1b or splicing regulatory (SRE) effectors (Pais et al., 2018; Huang et al., 2020; Derevnina et al., 2021).

In this study, we also reported that Rpi-amr3 directly interacts with AVRamr3 and other recognized AVRamr3 homologs from different *Phytophthora* species. Surprisingly, the direct interaction has not led to accelerated evolution of Avramr3 to evade detection, as we also observed for Rpi-amr1 and AVRamr1 (Lin et al., 2020a; Witek et al., 2021). This could predispose *Rpi-amr3* to function in different plant species.

Thus, *Rpi-amr3* could be deployed in Solanaceae crops like potato, tomato and tobacco against multiple *Phytophthora* diseases. However, interfamily transfer of *NLR* genes remains a challenge if *NLR* genes show “restricted taxonomic functionality” (Tai et al., 1999). The paired *NLR* genes *RPS4*/*RRS1* from Brassicaceae (*Arabidopsis*) can nevertheless function in other plant families like Solanaceae (tomato) and Cucurbitaceae (cucumber) against different bacterial and fungal diseases (Narusaka et al., 2013). Here, we found any of the NRC2, NRC3 or NRC4 proteins are required for Rpi-amr3 to execute its function. Thus, in the plant families which lacking NRC homologs, such as tropical tree crops susceptible to *P. palmivora* and *P. megakary*a, co-delivery of *Rpi-amr3* and *NRC* genes might be required to defeat these *Phytophthora* diseases.

“Non-host” resistance is durable. *S. americanum* and *S. nigrum* are thought to be non-host plants of *P. infestans*, although susceptible accessions of both species have been found using DLAs. This opens the opportunity to dissect their “non-host” resistance. By using AVRamr3 as a probe, we found Rpi-amr3 is widely distributed in *S. americanum* and *S. nigrum* species (Witek et al., 2021) (Fig. 6). We noticed that PITG_21190 (AVRamr3) triggers HR in many *S. nigrum* accessions in a large-scale effector screening study (Dong, 2016), consistent with our findings (Fig. 6). Furthermore, we cloned four *Rpi-nig3* genes from two *S. nigrum* accessions. All the four Rpi-nig3 homologs recognize AVRamr3 in our co-expression assays (Fig. 6), although their resistance to late blight needs to be evaluated individually. The wide distribution of *Rpi-amr3* and *Rpi-amr1* suggests that these two *R* genes, perhaps with other *R* genes and the NRC network in *S. americanum* and *S. nigrum*, underpin their “non-host” resistance against potato late blight. The identification of AVRamr3 and AVRamr1 can also help to explore other novel resistance genes from *S. americanum* and *S. nigrum*.

In summary, this study reveals that *Rpi-amr3* is a conserved and broad-spectrum *R* gene from *S. americanum* and its relatives. The recognition of the conserved AVRamr3 effectors leads to resistance against several different *Phytophthora* pathogens. This finding shows great potential for resistance breeding in many crop plants against different *Phytophthora* diseases.

## Acknowledgements

This research was financed from BBSRC grant BB/P021646/1 and the Gatsby Charitable Foundation. We thank TSL transformation team (Matthew Smoker and Jodie Taylor), SynBio team (Mark Youles) and horticultural team (Sara Perkins, Justine Smith, Lesley Phillips and Catherine Taylor) for their support. We thank Experimental Garden and Genebank of Radboud University, Nijmegen, The Netherlands, IPK Gatersleben, Germany and Sandra Knapp (Natural History Museum, London, UK) for access to *S. americanum* and *S. nigrum* genetic diversity. We thank He Meng and Lirui Cheng from CAAS for kindly sharing the *Phytophthora parasitica* isolates R0 and R1, and Franck Panabières from INRA for kindly sharing the *Phytophthora parasitica* isolates 310, 329, 666 and 721. We thank Joe Win from TSL for maintaining the *P. palmivora* strains. We thank Paul Birch and colleagues at James Hutton Institute for making available clones of some of the effectors that were tested for AVRamr3 function.

## Author contributions

X.L. and J.D.G.J. designed the study. X.L., A.C.O.A., R.H., K.W., H.S.K., T.S., C.-H.W. and H.A. performed the experiments. X.L., A.C.O.A., R.H. and K.W. analysed the data. X.L. and J.D.G.J. wrote the manuscript with input from all authors. S.K. and V.G.A.A.V. contributed resources. All authors approved the manuscript.

## Conflict of interest

K. W. and J.D.G.J. are named inventors on a patent application (PCT/US2016/031119) pertaining to *Rpi-amr3* that was filed by the 2Blades Foundation on behalf of the Sainsbury Laboratory. The other authors declare no competing interests.

## Supplementary files

**Figure S1:**
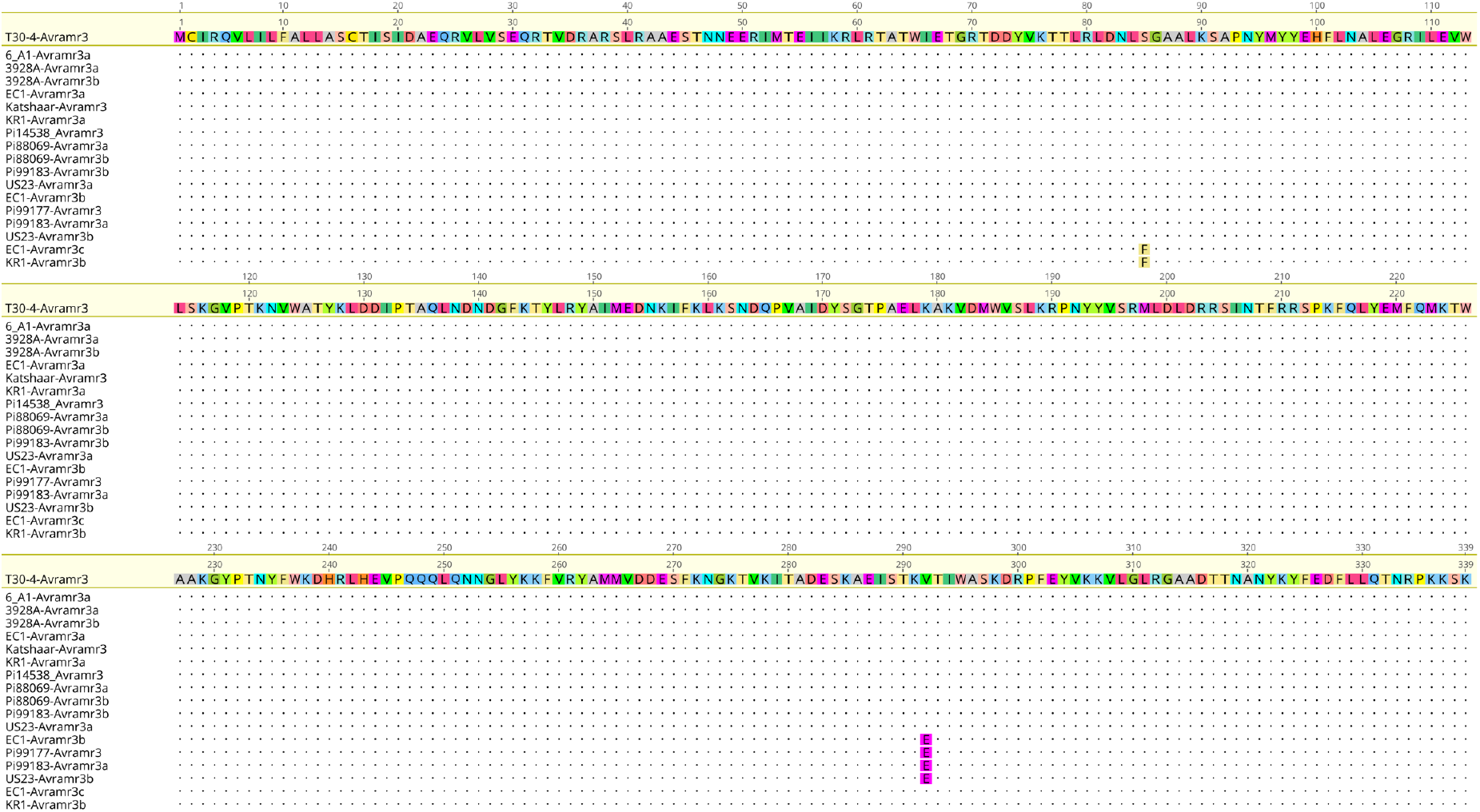
Protein alignment reveals strong conservation of *P. infestans* AVRamr3 alleles and paralogs

**Figure S2:**
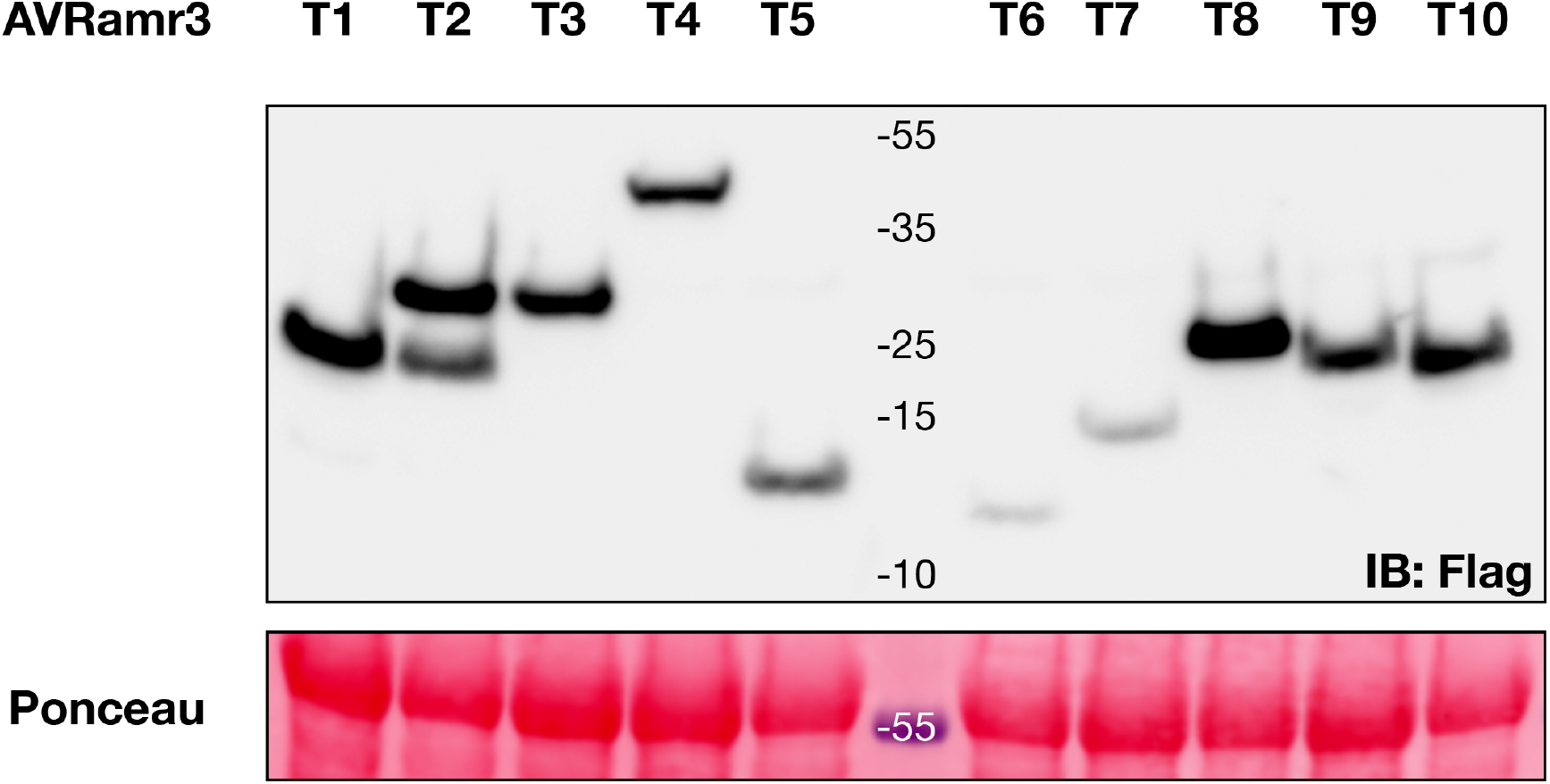
Western blot for AVRamr3 truncations with C-terminus HIS-FLAG tag.

**Figure S3:**
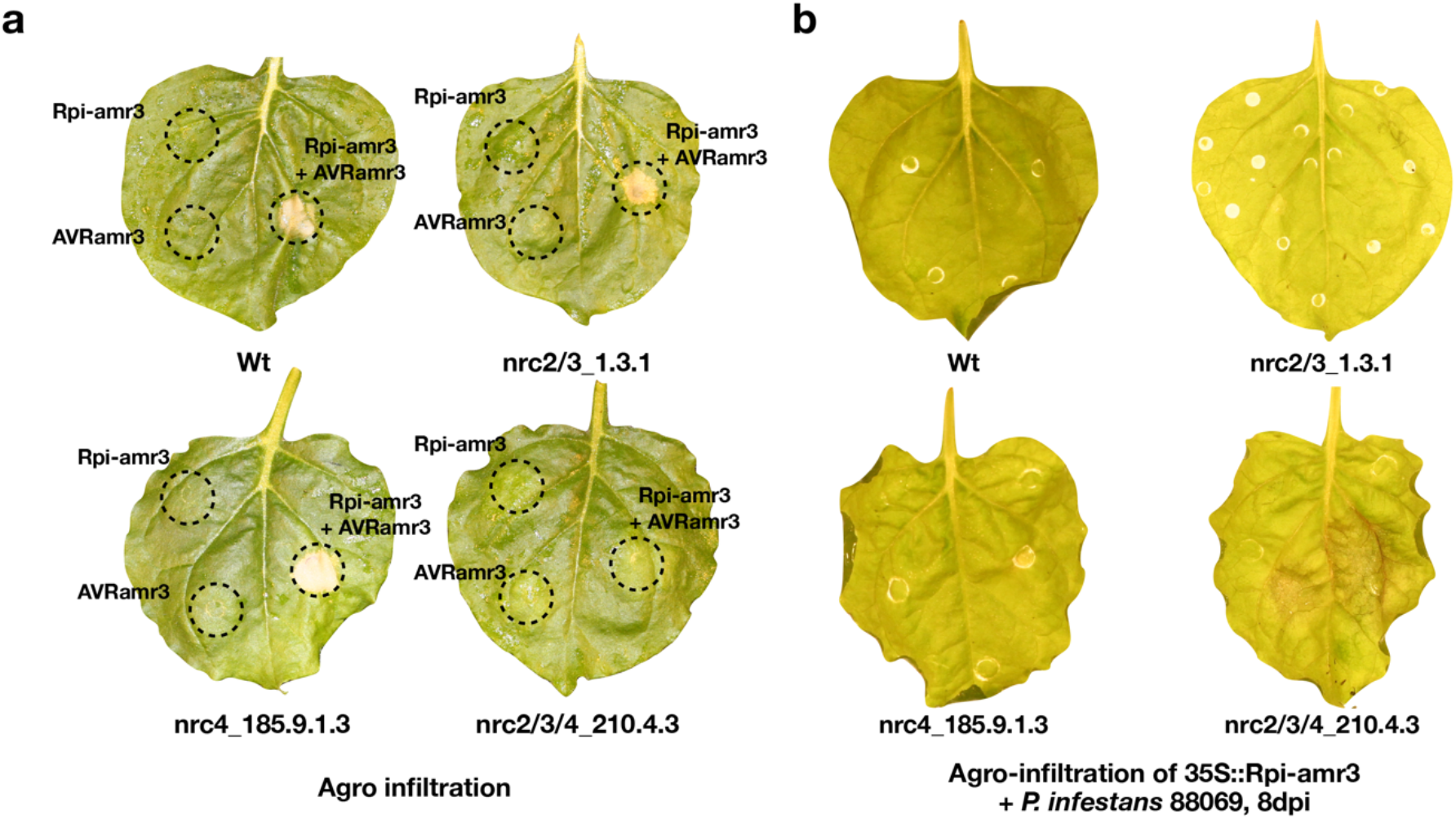
Rpi-amr3 is NRC2, NRC3 or NRC4 dependent. (a) HR phenotype after expressing Rpi-amr3, AVRamr3 and Rpi-amr3+AVRamr3 in wild type, nrc2/3_1.3.1, nrc4_185.9.1.3 and nrc2/3/4_210.4.3 knockout *N. benthamiana* lines. (b) Detached leaf assay (DLA) after *Rpi-amr3* transient expression in wild type, nrc2/3_1.3.1, nrc4_185.9.1.3 and nrc2/3/4_210.4.3 knockout *N. benthamiana*. 500 zoopores of *P. infestans* 88069 were used one day after Rpi-amr3 transient expression by agro-infiltration. The photos were taken eight days after inoculation.

**Figure S4:**
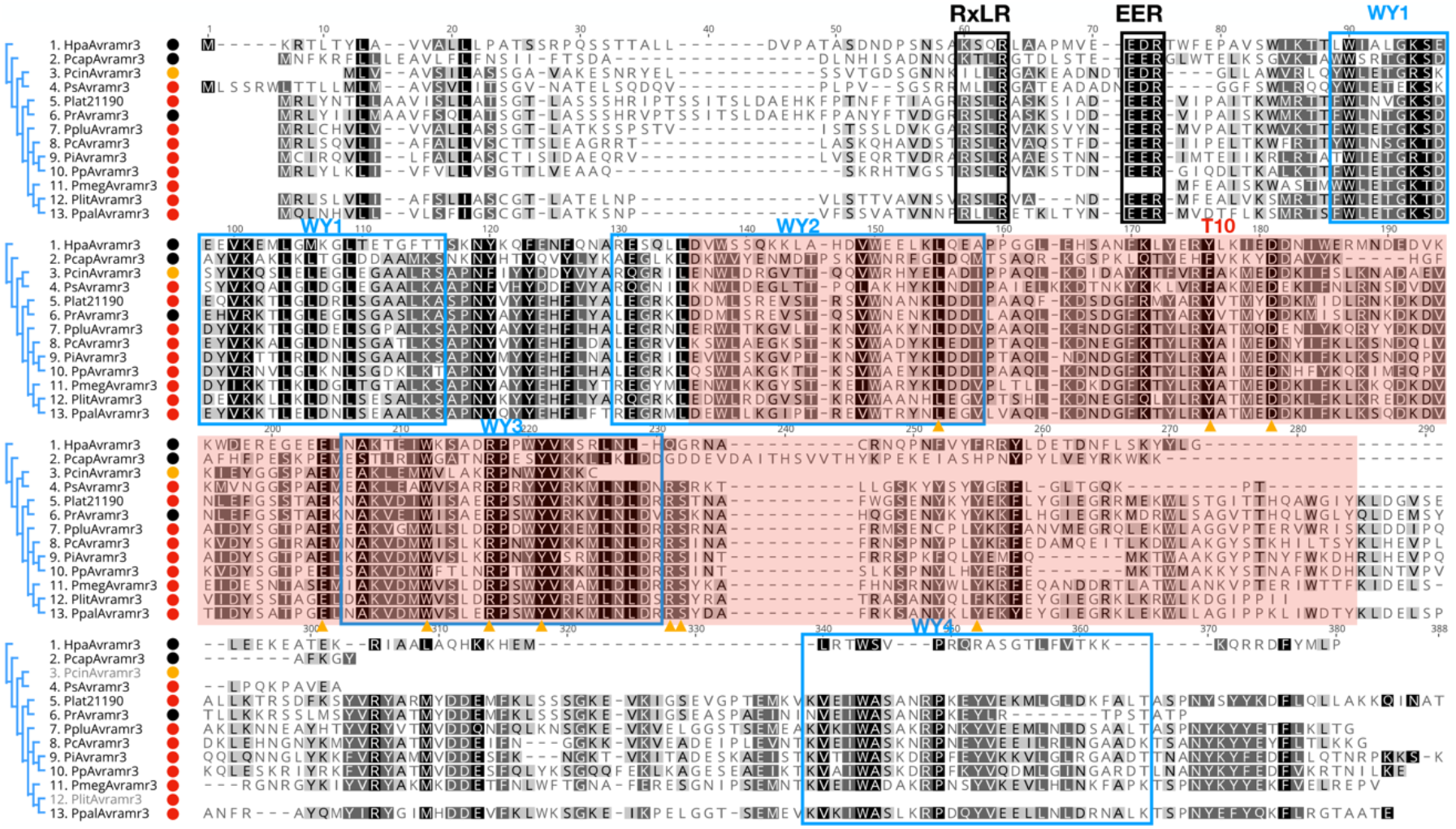
Protein alignment of AVRamr3 homologs from different *Phytophthora* genomes, including *Phytophthora infestans* (Pi), *Phytophthora parasitica* (Pp), *Phytophthora cactorum* (Pc), *Phytophthora palmivora* (Ppal), *Phytophthora megakarya* (Pmeg), *Phytophthora litchi* (Plit), *Phytophthora sojae* (Ps), *Phytophthora lateralis* (Plat), *Phytophthora pluvialis* (Pplu), *Phytophthora ramorum* (Pr), *P. cinnamomi* (Pcin), *P. capsica* (Pcap) and *Hyaloperonospora arabidopsidis* (Hpa). The circles after the name are their recognition specificity by Rpi-amr3 in the HR assay, red: HR; black: no HR; yellow: weak HR. The RXLR and EER motifs are marked by black boxes. The predicted WY motifs are marked by blue boxes. The conserved amino acids which were selected for mutagenesis (see Figure S5) are marked by yellow arrows.

**Figure S5:**
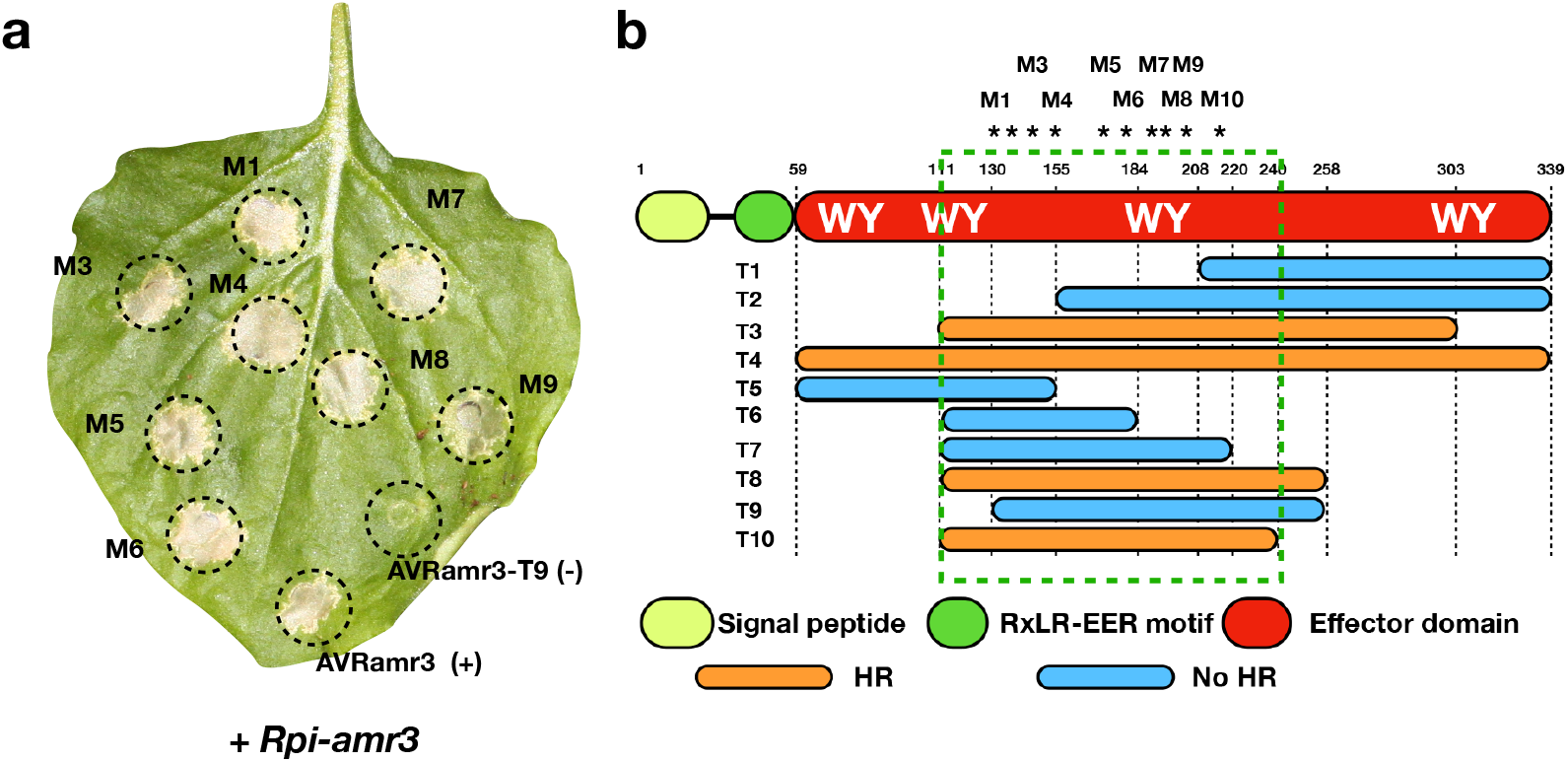
Mutagenesis of AVRamr3 from *Phytophthora infestans*. **(a)** Eight *Avramr3* mutants were generated by PCR and cloned into over-expression vector with 35S promoter. All of them induce HR when co-expressed with Rpi-amr3. An AVRamr3 truncation T9 was used as negative control, full-length AVRamr3 was used as positive control. **(b)** The position of each mutation is marked by asterisk, and correspond to those amino acids marked by yellow arrows in Fig S4. All the mutants are in the T10 region and are conserved among different AVRamr3 homologs from other *Phytophthora* species (see Figure S4).

**Figure S6:**
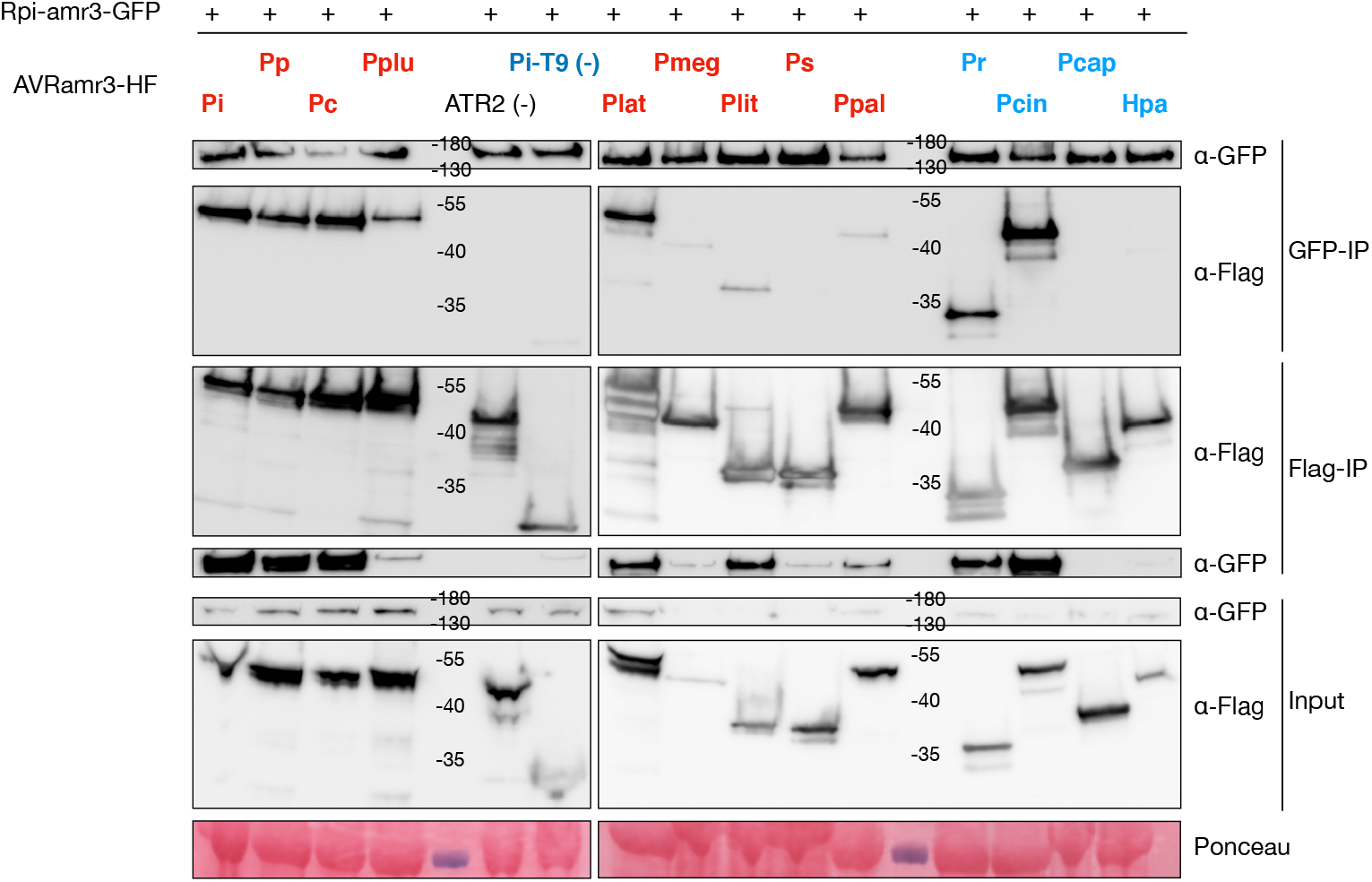
Co-IP of Rpi-amr3 and AVRamr3 homologs. Rpi-amr3 is tagged with C-terminal GFP, and all AVRamr3 homologs are fused with a C-terminal HIS-FLAG tag. Bi-directional Co-IPs were performed by GFP-IP or Flag-IP individually then incubated with Flag-HRP or GFP-HRP antibody. ATR2 and AVRamr3 truncation T9 were used as negative controls.

**Figure S7:**
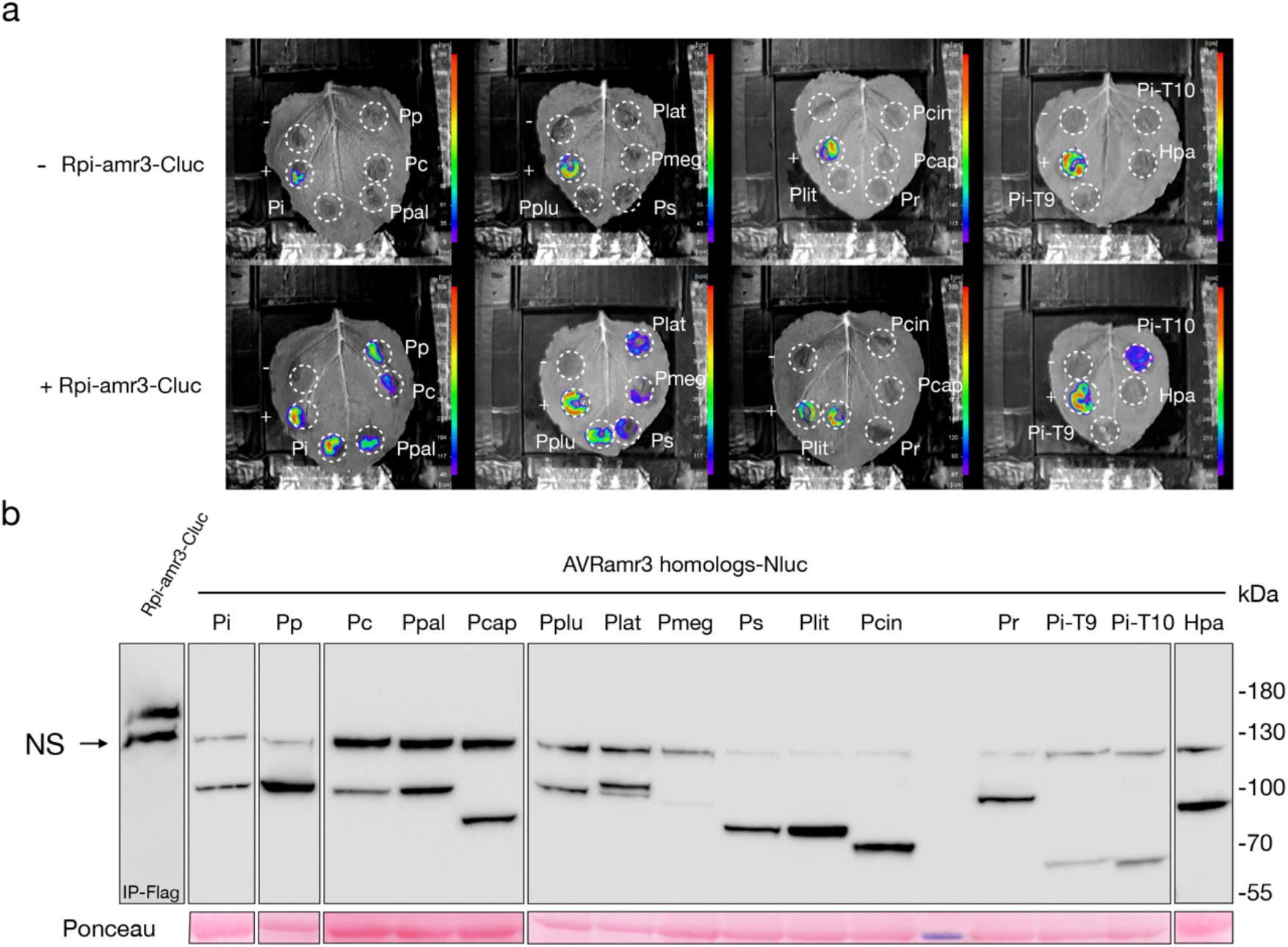
Split luciferase for Rpi-amr3 and AVRamr3 homologs from different *Phytophthora* species. Rpi-amr3 was fused with Flag-Cluc and AVRamr3 homologs were fused with Flag-Nluc. The experiment was performed in nrc2/3/4_210.4.3 knockout *Nicotiana benthamiana* to abolish HR. **(a)**. Expressing the AVRamr3::Flag-Nluc homologs alone does not show luciferase signal; Co-expression of Rpi-amr3::Flag-Cluc with AVRamr3::Flag-Nluc homologs can induce luciferase signal, specifically, *P. infestans* (Pi), *P. parasitica* (Pp), *P. cactorum* (Pc), *P. palmivora* (Ppal), *P. megakarya* (Pmeg), *P. litchi* (Plit), *P. sojae* (Ps), *P. lateralis* (Plat) and *P. pluvialis* (Pplu) interact with Rpi-amr3::Flag-Cluc; AVRamr3 homologs from *P. ramorum* (Pr), *P. cinnamomi* (Pcin), *P. capsici* (Pcap) and *Hyaloperonospora arabidopsidis* (Hpa) do not interact with Rpi-amr3 in this assay. **(b)**. Western blot by FLAG antibody was performed to confirm the expression of all proteins.

**Figure S8:**
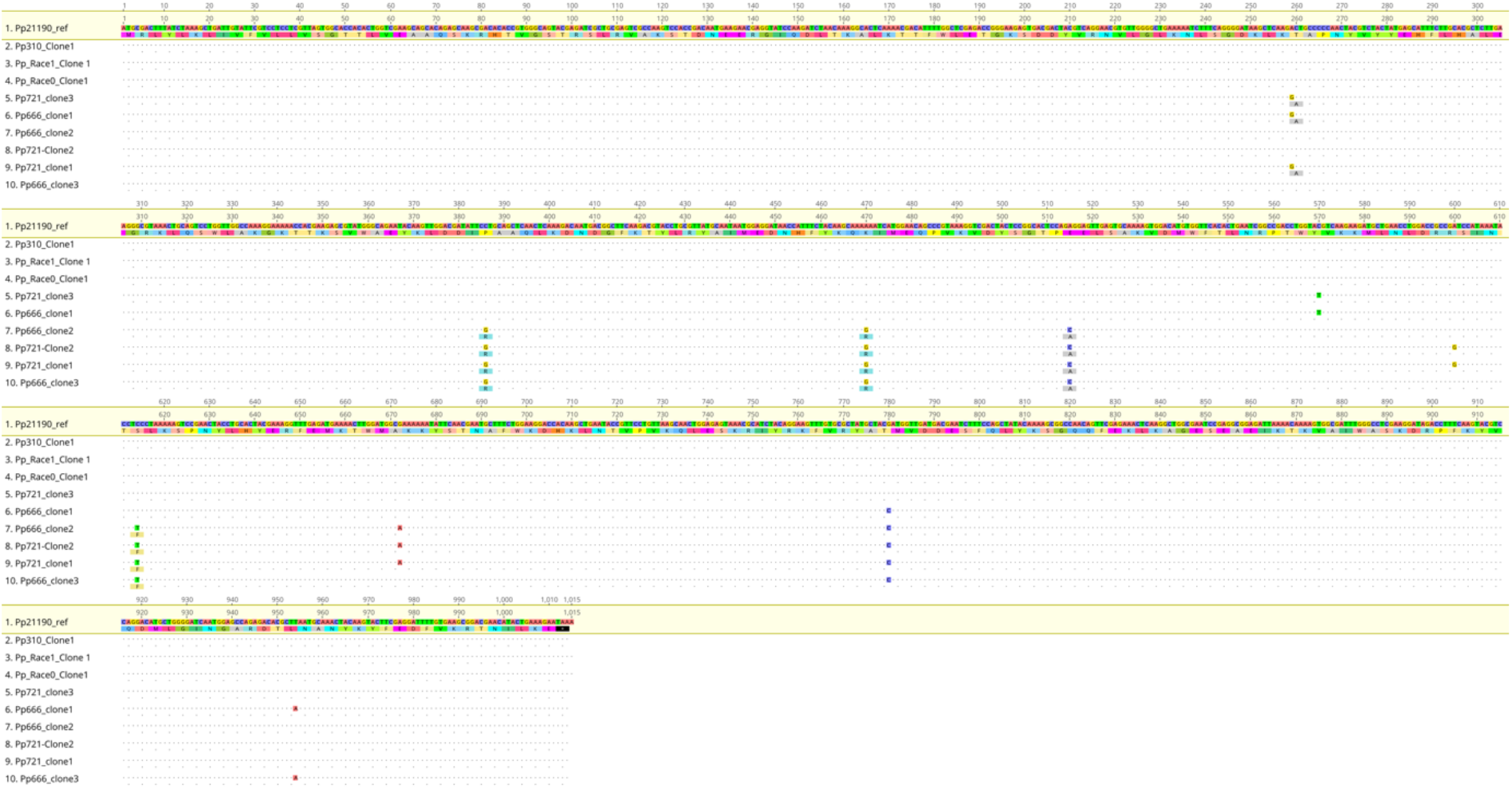
DNA alignment of *Avramr3* homologs from different *Phytophthora parasitica* isolates. The polymorphic DNA and amino acids are highlighted.

**Figure S9:**
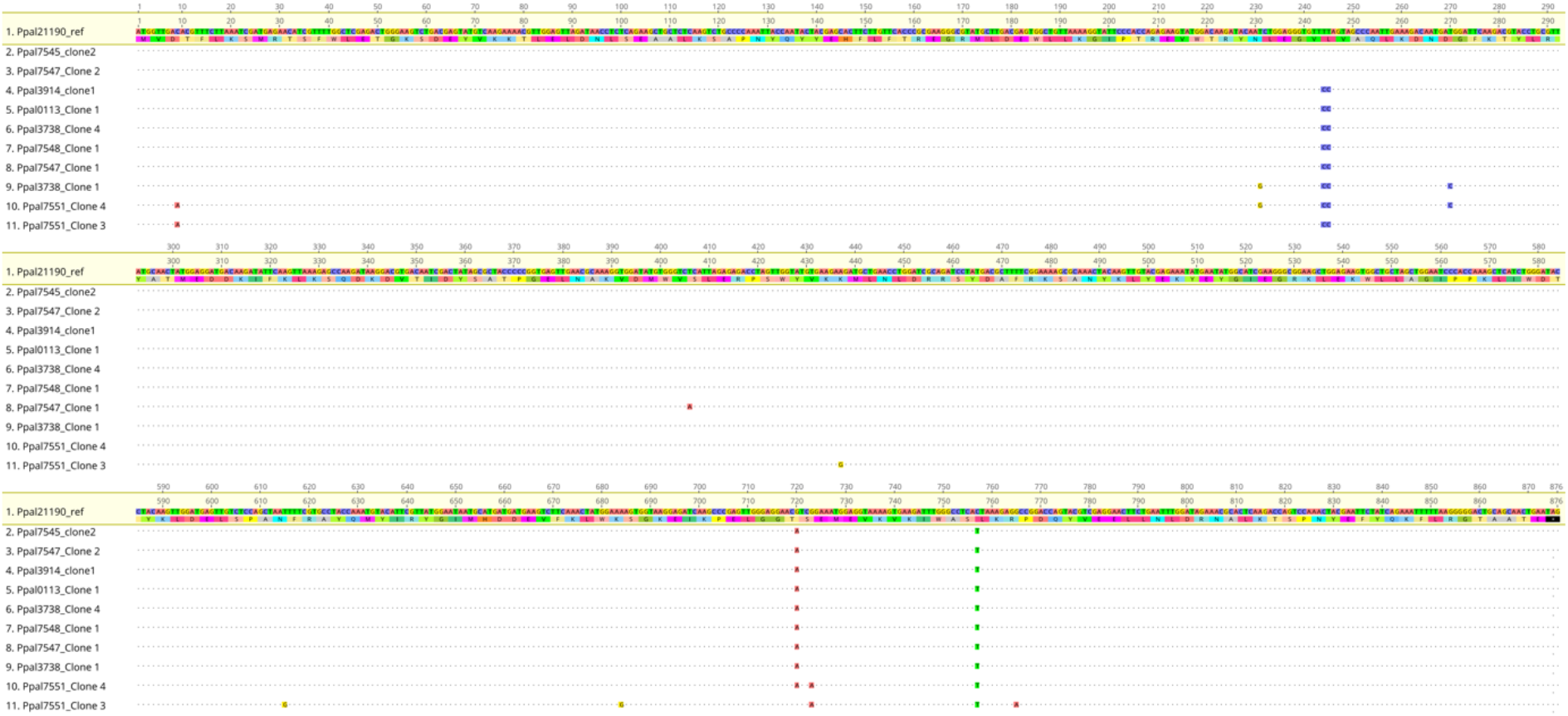
DNA alignment of *Avramr3* homologs from different *Phytophthora palmivora* isolates. The polymorphic DNA and amino acids are highlighted.

**Figure S10:**
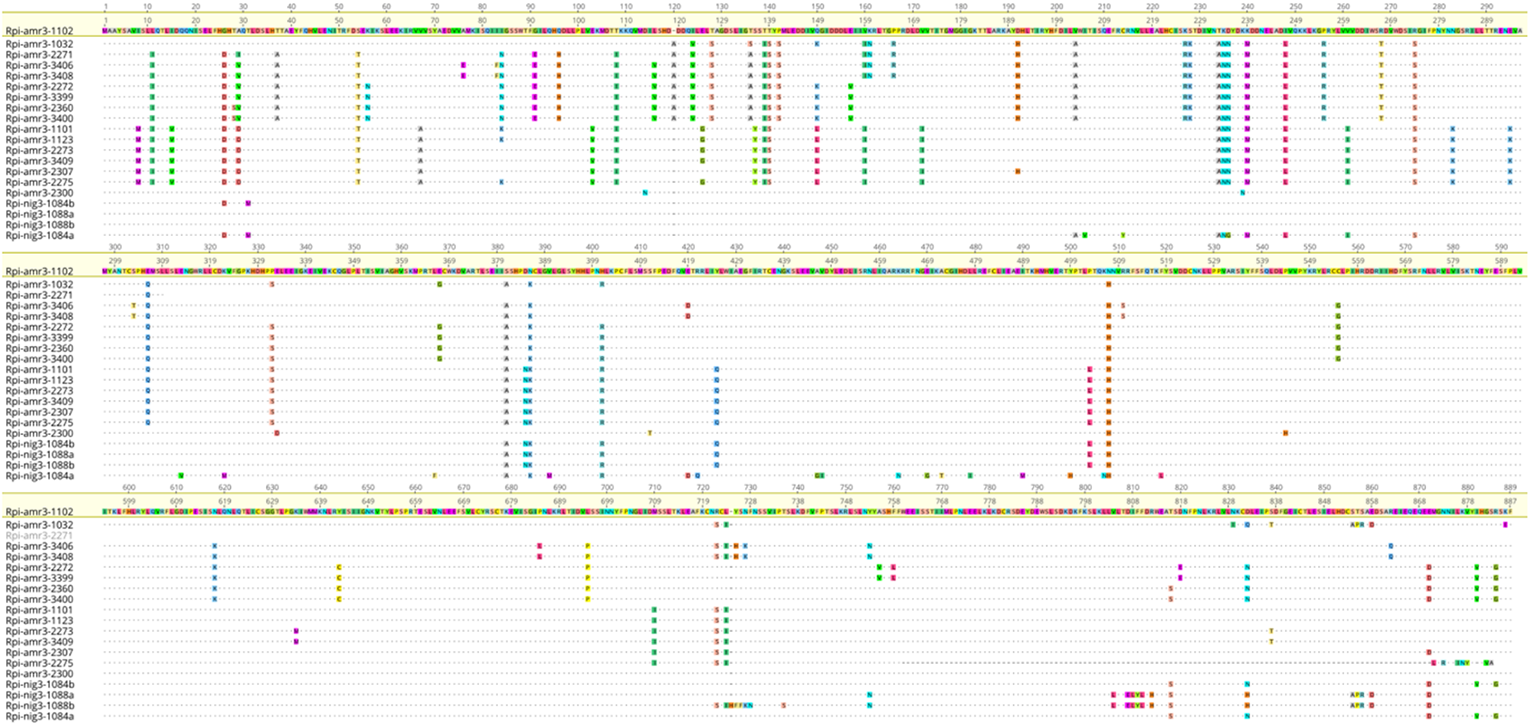
Protein alignment of Rpi-amr3 and Rpi-nig3 homologs from *Solanum americanum* and *Solanum nigrum* accessions. The Rpi-amr3 was used as reference, all the polymorphic amino acids are highlighted.

