## Supplementary material for "*Rpi-amr3* confers resistance to multiple *Phytophthora* species by recognizing a conserved RXLR effector": Material and Methods

**Materials and Methods**

**RXLR effector libraries**

The RXLR effector libraries were described previously (Rietman, 2011; Lin et al., 2020). *Rpi-amr3* with its native promoter and terminator (Witek et al., 2016) was co-expressed with individual effectors in *N. benthamiana* by agro-infiltration. And the HR phenotype were scored three days after the agro-infiltration.

**Plant materials**

The plant materials used in this study are listed in Table S1, the *Nicotiana benthamiana* NRC2/3, NRC4 and NRC2/3/4 knockout lines are described previously (Adachi et al., 2019; Wu et al., 2020; Witek et al., 2021), T2 *N. benthamiana-Rpi-amr3* transgenic lines were generated, full-length *Rpi-amr3* gene with its native promoter and terminator was cloned into a binary vector, and used for the *N. benthamiana* transformation (Witek et al., 2016), two homozygous T2 lines *Rpi-amr3*#13.3 and *Rpi-amr3*#16.5 were selected by detached leaves assays (DLA), 15 T2 plants of each *Rpi-amr3*#13.3 and *Rpi-amr3*#16.5 lines were tested, and all are resistant to *P. infestans* isolate 88069. The wild type, knockout and transgenic *N. benthamiana* were propagated in a glasshouse, for the experiments, the plants were grown in a controlled environment room (CER) with 22 °C, 45-65% humidity and 16 hours photoperiod.

The *S. americanum* and *S. nigrum* accessions were collected from different seed banks, the seeds were sowed and grown in containment glasshouse for agro-infiltration experiments.

**Pathogens and disease test**

The pathogens used in this study are listed in Table S1. *Phytophthora infestans* isolate 88069 was used for the *P. infestans* disease test, they were propagated and maintained on rye sucrose agar (RSA) medium in a 18°C incubator, ice cold water was used to induce zoospores from 7-14 days old plates. The plate then incubated at 4 °C for an hour then the zoospores suspension was collected and used for detached leave assay (500 zoospores/droplet). Both *Phytophthora parasitica* and *Phytophthora palmivora* isolates were propagated and maintained in V8 plates in a 25°C incubator. To produce the zoospore from *P. palmivora*, 7-10 days old plates were flooded with 4 °C water and incubated at 4 °C for 1h, then moved to room temperature for another 1h, the released zoospores were counted by a hemocytometer, 30,000/mL - 50,000/mL zoospores were used for the root inoculation, 1 mL zoospore suspension were added to the root. For *P. parasitica*, 10 days old plate was flooded by 0.1% KNO_3_ solution, incubated in 25°C for 2 days in dark, then incubate in 4 °C for 1 hour and 25 °C for another 1h to release the zoospores. 30,000/mL - 50,000/mL zoospores were used for the root inoculation, 1 mL zoospore suspension were added to the root. 3 - 4 weeks wild-type or *Rpi-amr3* transgenic *N. benthamiana* plants were used in the disease test, they were grown in CER, and the inoculated plants were grown in a Sanyo cabinet with 25 °C and 16 h photoperiod, the root inoculation experiment takes 4-7 days, the scoring was taken when the wild-type *N. benthamiana* plants are infected completely.

**Genomic and sequence analysis**

The *Phytophthora* genomes used in this study are listed in Table S1 . The genomes were imported into Geneious R10 (Kearse et al., 2012), and local BLAST databases were generated. The *Avramr3* homologs containing contigs were identified by BLAST, then *Avramr3* loci were extracted with flanking sequences. All the *Avramr3* loci were re-annotated by gene prediction tool in EumicobeDB (<http://www.eumicrobedb.org/eumicrobedb/gene_predict.php>) (Panda et al., 2018). Then the GenBank (.gb) file was exported and visualized by Clinker (https://github.com/gamcil/clinker) (Gilchrist and Chooi, 2021). All other sequence were analysed in Geneious R10. All sequence alignments were performed by MAFFT (Katoh and Standley, 2013). The phylogenetic tree was generated by IQ-tree (v1.6.12) (Minh et al., 2020), 546 protein models were tested and JTT+G4 was selected as the best-fit model, 1000 samples were generated for the ultrafast bootstrap analysis.

**Molecular cloning and constructs used in this study**

All constructs and primers used in this study are listed in Table S1. In brief, the *Rpi-amr3* CDS were cloned into golden gate level 0 entry vector pICSL01005, then fused with C-HA (pICSL50009), C-GFP (pICSL50008) or C-Flag-Nluc (pICSL500047) tags and recombined into level 1 vector pICSL86977OD with 35S promoter and Ocs Terminator. All the AVRamr3 homologs from different *Phytophthora* species, were synthesized based on their reference genomes, the signal peptides were removed, and BsaI and BpiI sites were domesticated to facilitate the golden gate cloning. All the *Avramr3* homologs, truncated *Avramr3* and *Avramr3* point mutants were cloned into level 0 entry vector pICSL01005, then fused with C-HIS-FLAG (pICSL50001) or C-Flag-Cluc (pICSL500048) tags. The *Rpi-amr3/Rpi-nig3* homologs from *S. americanum* and *S. nigrum* were amplified by nested PCR and cloned into level 1 vector pICSL86922OD containing a 35S promoter and Ocs Terminator.

For generating the *Rpi-amr3* transgenic *N. benthamiana* lines, *Rpi-amr3* with its native promoter and terminator were cloned into USER vector pICSLUS0001OD (Witek et al., 2016), and shuffled into *Agrobacterium* strain AGL1 for plant transformation. The *N. benthamiana* plants were propagated in a glasshouse, two homozygous T2 lines were selected for the *Phytophthora* disease test.

**Agro-infiltration**

All the over-expression constructs were shuffled into *Agrobacterium* strain GV3101-pMP90, and they are stored in -80 °C freezer with 20% glycerol. The *Agrobacterium* were streaked out on solid L medium with antibiotics and incubated at 28 °C for two days, then the *Agrobacterium* were re-suspended into infiltration buffer (MgCl_2_-MES, 10mM MgCl_2_ and 10mM MES, pH 5.6) with 1 mM acetosyringone and used for agro-infiltration, OD_600_ = 0.5.

**Western blot and co-immunoprecipitation**

The Western blot and co-immunoprecipitation protocols were described previously (Guo et al., 2020). In brief, 35S::Rpi-amr3::HA or 35S::Rpi-amr3::GFP, 35S:: AVRamr3::HIS-FLAG or other AVRamr3 homologs with C-HIS-FLAG tag were transiently co-expressed in *N. benthamiana* NRC2/3/4 knockout line, OD_600_ = 0.5. The leaves were sampled at 3 dpi and total protein were extracted by GTAN buffer for co-immunoprecipitation. EZview^TM^ Red Anti-HA Affinity Gel (Sigma-Aldrich, Cat: E6779), Anti-Flag^®^ M2 affinity gel (Sigma-Aldrich, Cat: A2220), GFP-Trap Agarose (ChromoTek, Planegg-Martinsried, Germany) were used for the immunoprecipitation, HRP conjugated HA antibodies (Sigma-Aldrich, Cat: H6533), HRP conjugated anti-FLAG antibodies (Sigma, Cat: A8592) and HRP conjugated anti-GFP antibodies (Santa Cruzm Cat: sc-9996HRP) were used for the Western blot. NuPage 4-12% Bis-Tris protein gels (ThermoFisher, Cat: NP0302BOX) and MOPs SDS Running Buffer (ThermoFisher, Cat: NP0001) were used for separating the protein.

**Split-luciferase assay**

The split-luciferase system was described previously (Chen et al., 2008). *Rpi-amr3* was fused with 1x Flag::C-luciferase (Cluc) tag and *Avramr3* homologs were fused with 1x Flag::N-luciferase (Nluc), the Rpi-amr3::Cluc and AVRamr3::Nluc were transiently co-expressed in *N. benthamiana* NRC2/3/4 knockout line by agro-infiltration. Three days after infiltration, 0.4 mM luciferin on 100mM sodium citrate buffer (pH 5.6) was infiltrated into the leaves, then the leaves were detached for imaging (NightOWL II LB 983 in Vivo Imaging System with WinLight^32^ Software, BERTHOLD TECHNOLOGIES GmbH & Co KG, Germany). Two leaves were used for each experiment, and three independent biological repeats were performed. Western blots with HRP-FLAG antibodies were used for detecting the presence of the fused protein.
